# Use of ANCHOR viruses to validate radiative, physical and chemical decontamination systems in operational conditions

**DOI:** 10.1101/2025.04.15.648913

**Authors:** Franck Gallardo, Valentine Poirier, Charlotte Quentin-Froignant, Thomas Figueroa, Romain Volmer, Morgane Mousnier, Olivier Authier, Dani Lainisalo, Pertti Lainisalo, Tero Ingelius, Marc Grandadam, Elie Marcheteau

**Author notes:** Corresponding authors (F. Gallardo), (E. Marcheteau).

## Abstract

ANCHOR tagged viruses provide a high-performance platform for the rapid evaluation of disinfection methods across chemical, radiative, and physical modalities. This study highlights the robustness of Vaccinia virus (VACV) and human Adenovirus type 5 (hAdV5) as environmental persistence models, demonstrating their suitability for testing disinfection protocols. ANCHOR imaging capability allows real-time visualization of disinfection efficacy, enabling rapid calibration and cost-effective validation of new disinfection systems. This technology provides essential support for the development and optimization of disinfection solutions in public areas up to the improvement of safety protocols in high-containment laboratories. The increasing number of BSL-4 laboratories worldwide underscores the critical needs for efficient and reliable decontamination systems and methods to test them in operational conditions to ensure biohazard safety.

**Highlights:**

1. ANCHOR-tagged viruses provide a rapid and cost-effective platform for evaluating disinfection methods across chemical, physical, and radiative modalities.
2. Human Adenovirus type 5 (hAdV5) and Vaccinia virus (VACV) serve as robust environmental persistence models for testing viral decontamination protocols.
3. The use of ANCHOR viruses enables rapid real-time visualization of disinfection efficacy, improving safety validation in high-containment laboratories and public health applications.

## Introduction

Emerging or re-emerging virus outbreaks can have a tremendous impact on the general population in terms of public health and economic shutdown, as we have seen recently during the Covid19 pandemic^1^. The Covid19 crisis raised the need of new efficient and high throughput disinfection strategies, both at individual and collective scales. In another field, by 2023, there are 51 biosafety level 4 (BSL-4) laboratories in operation around the world, with a further 18 facilities currently scheduled or under construction stage^2^. This wave of interest in high-containment laboratories began in 2001, following the anthrax attacks in the USA, but more recently the SARS-CoV-1 and Covid19 epidemics have led the international community to acquire this type of laboratory. This poses a number of concerns, particularly for the surveillance of experimental programs when laboratories are hosted by countries with no or poor biohazards regulation control^3^.

Current decontamination methods for viral inactivation lays on the sensitivity of viral components to either chemical, physical or radiative sources. Chemical disinfection may involve oxidizing or fixating agents (NaOCl, H_2_0_2_, ClO_2_, etc.), alcohol-based agents and aldehydes whereas physical decontamination relies on heat, filtration and desiccation for example. Radiative decontamination can use several sources but the ultraviolet type C (UV-C) is one of the most used in infectious context^4^. The impact of UV energy on the virus, mainly on the virus genome, is the formation of covalent bonds between two adjacent nucleotides making viral replication difficult or impossible. It is interesting to note that RNA viruses are theoretically less sensitive to UV than DNA viruses, as uracil dimer is more difficult to form than pyrimidine dimer^5^. Remarkably, Adenoviruses, a family of dsDNA viruses triggering several pathologies in human and animals, are highly resistant to UV-C because they can take advantage of the cell repair mechanisms of nucleic acid alterations. This resistance, even though they are DNA viruses, reinforces the idea of using a labelled Adv as a model for rapid testing of many irradiation conditions^6^. Classical techniques to investigate remaining infectious viral particles involves titration by either plaque reduction assay scored in plaque forming unit (PFU/mL) or tissue culture infectious dose 50% (TCID50/mL)^7^. These techniques are extremely precise but need large amount of cell culture conditions and are time consuming with massive hands-on investment. A technology that will allow to measure rapidly in miniaturized condition the efficiency of a disinfection process by providing a direct readout could be of help to reduce time to result and to lower the amount of infectious wastes generated. To this end, we have developed various viral models of DNA-tagged fluorescent viruses in our laboratory, using our ANCHOR system^8–10^, and use these viruses in high content microscopy in 96 well plates to visualize and quantify infectious virus presence scoring the virus as fluorescent forming units (ffu/ml). By quantifying only infectious viral particles, this system not only enables us to rapidly monitor infection, but also offers us the possibility of using these models for viral persistence studies on a wide range of materials. We have adapted this system to the evaluation of all types of disinfection, whether radiative, chemical or physical by selecting the most relevant viruses based on their natural resistance. Our rapid implementation means we can intervene at a very early stage in the development of disinfection systems, but we can also intervene on an *ad hoc* basis to check the quality of the systems produced. The viruses used are also of low biosafety level, limiting the risk to operators or the community. Our experience of the covid19 pandemic has shown us that the quality of components in disinfection systems can be highly variable, despite regulatory certification, and disinfection efficiency can vary depending on the nature of the virus tested and the nature of the contaminated material. It is therefore of high importance, even for a disinfection method that has been normatively accredited, to verify its efficiency at the point of use in operational conditions when possible. This is of high concern for BSL-3 and BSL-4 laboratories, where biohazard contamination can lead to tremendous impact in the environment and in the general population^11–13^.

In this study, we used our virus collection to test radiative, physical and chemical disinfection methods in operational environments. We were able to independently verify the efficiency of the procedure used to disinfect pressurized suits used in French army BSL-4 labs, highlighting points of vigilance that were not previously known. To do this, we used a VACV model virus tagged with the ANCHOR system for rapid analysis of the various conditions tested. This poxvirus has the advantage of mimicking the behavior of other poxviruses of civil and military interest, such as monkeypox or smallpox. Indeed, their stability in the environment makes them a good model, given the category of viruses used in the BSL-4 of interest. VACV ANCHOR was also used to contaminate the showering room for BSL-4 entry/exit and measure disinfection in operational conditions during nebulization.

## Results and discussion

### Use of autofluorescent ANCHOR viruses to measure persistence of infectivity of several viral strains

We have previously shown that infection and replication capacities of ANCHOR tagged viruses are not modified *in vitro*^8,14^. However, we do not know if these engineered viruses will present a defect in viral stability due to the insertion of the ANCHOR system and accumulation of OR protein onto the tagged genome. We choose as diluent the water as this can mimic the effluent of BSL-4 showers following rinsing. To verify this, we spiked water samples with an enveloped virus considered to be resistant in the environment (poxvirus, VACV ANCHOR), a non-enveloped virus considered to be resistant in the environment (Adenovirus, hAdv5 ANCHOR) and an enveloped virus considered to be sensitive in the environment (Herpesvirus, hCMV ANCHOR) at the indicated concentration at Day 0 (1.10^7^ ffu/mL for hAdv5 ANCHOR, 3,2.10^5^ ffu/mL for VACV ANCHOR and 1,5.10^7^ ffu/mL for hCMV ANCHOR). Transferring viruses from congelation medium to water can have an impact on their infection capacities. We therefore calculated the remaining infectious particles immediately after transferring viruses in water samples at Day 0. Compared to the targeted concentrations, we calculated an initial infectious virus concentration of 7,47.10^6^ (+/-0,43), 1,77.10^5^ (+/-0,09) and 2,44.10^6^ (+/-0,24) ffu/mL for hAdv5, VACV and hCMV respectively. Water transfer therefore triggered a 22, 45 and 82% drop in the number of infectious particles for hAdv, VACV and hCMV respectively. These results may underline the intrinsic resistance of non-enveloped viruses (hAdv) to environmental changes, notably osmotic pressure. For enveloped viruses, VACV was far more resistant than hCMV. Contaminated water samples were then incubated at either 4°C, room temperature (RT) or 37°C and aliquot were used to infect permissive cells and score remaining infectious virus particles at the indicated time. In line with existing literature^15^, we found that hCMV was very unstable in the contaminated sample under all conditions tested, with a massive (>50%) drop in infectivity level after 48 h.

After a week, only 10% of particles were still infectious. After two weeks, no infectious particle can be detected in the RT and 37°C conditions, whereas remaining infectious particles are maintained at 4°C up to three months (300 ffu/mL). For VACV and hAdv5, these viruses proved to be relatively stable over time at RT (around 20 days to lose 50% infectivity), but infectivity decreases rapidly at 37°C. However, at 4°C for 120 days, VACV had lost only 40% of its infectivity and hAdV5 60% (Figure 1). These data underline the fact that hAdV5 and VACV are good models for the evaluation of viral persistence in the environment and disinfection efficiency. Disinfection methods are vast and can be based on several principles to finally achieve virus inactivation, these methods can be radiative, physical or chemical.

**Figure 1.**
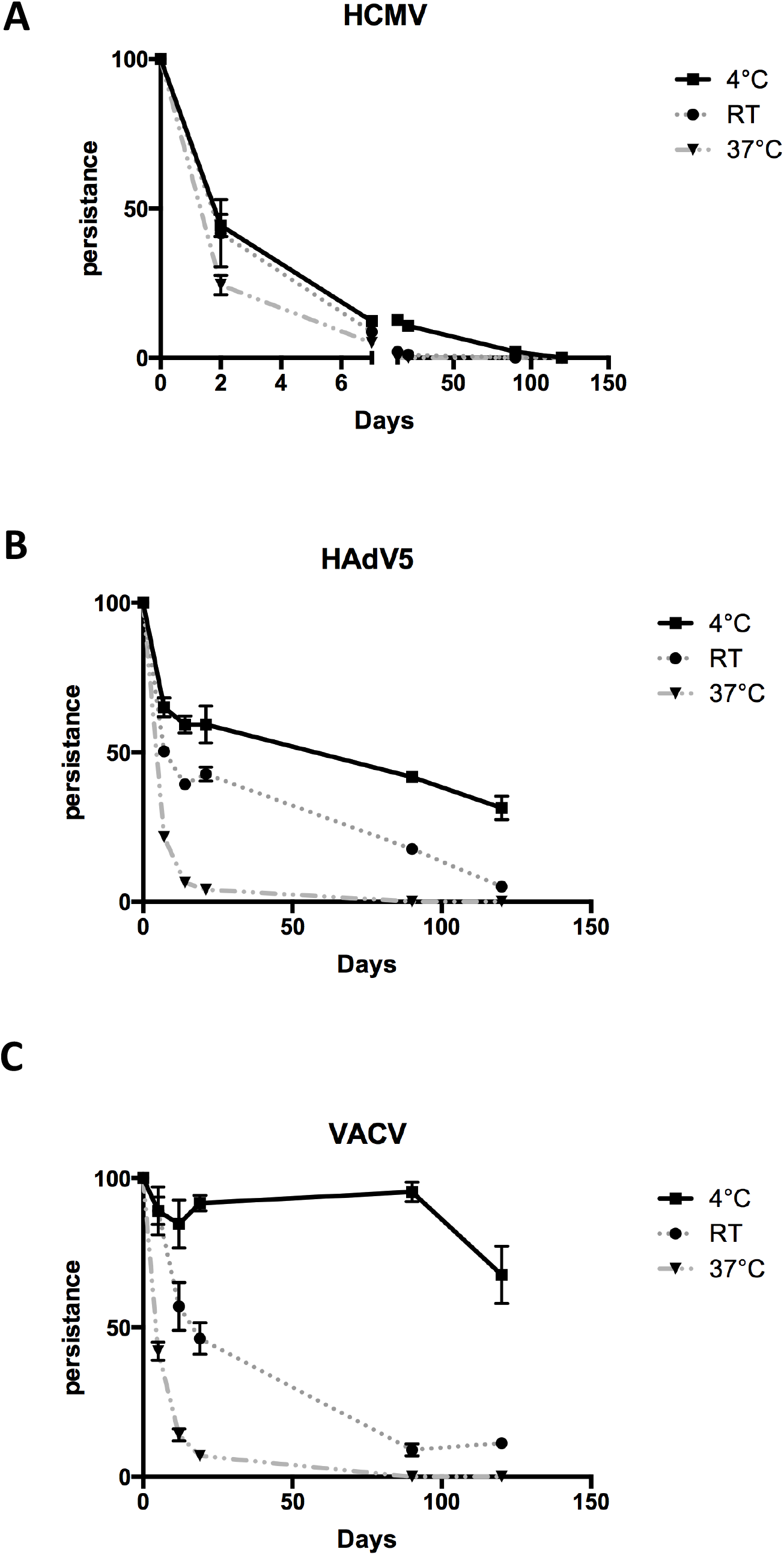
Evaluation of the impact of different temperature conditions on ANCHOR™ Viruses. **A:** hCMV ANCHOR™ persistence in water at temperatures of 4°C, 37°C and at room temperature. **B:** hAdV5 ANCHOR™ persistance in water at temperatures of 4°C, 37°C and at room temperature. **C:** hCMV ANCHOR™ persistance in water at temperatures of 4°C, 37°C and at room temperature. At each indicated timepoints number of infectious particles were calculated by infecting permissive cells, up to day120.

### Combining autofluorescent viruses and UV irradiation to develop and validate disinfection devices

The emergence of Covid19 and its rapid spread in the general population worldwide led to a strong effort to develop and validate, at affordable cost, home based disinfection devices specifically targeting viruses. *Adenoviridae* family shows a higher resistance to UV-C than *Coronaviridae*^6,16,17^. In fact, in the scientific literature, to achieve a virus inactivation of 4 Log10 (99,99%), it is necessary to reach 216 and 240 mJ/cm^2^ for AdV5 against 91 mJ/cm^2^ for SARS-CoV-2.

We started to work on the development of a disinfection box (e-Box, Clinit technologies) and used our autofluorescent hAdv5 ANCHOR virus as a test virus to investigate viral inactivation. The first prototypes we tested were 3D-printed and realized in house (Figure 2A). A known hAdv5 ANCHOR viral load is deposited on fabric or plastic samples, which are then dried and irradiated with 230 to 1400 mJ/cm^2^ UV-C for a 10-minute cycle. After irradiation, the residual viral particles are recovered, used to infect permissive cells overnight and scored by high content microscopy (Figure 2B). The first series of tests demonstrated an efficacy of up to 3 log10 (99,9%) disinfection on plastic samples and 1 log10 on textiles (90%) (Figure 2C). Similar results were obtained when the sample was placed directly under the LED ramp or off-centered. The poor inactivation on textile samples led our partner to increase the intensity of the LEDs to get closer to 4 log10 efficacy when producing the first series of e-Box. Therefore, the device was modified accordingly and a pre commercial production of 1000 boxes was fabricated with higher power. We repeated testing on this new improved device as shown on Figure 2C. The results were far from what was expected, as the device had difficulty achieving 95% decontamination depending on the position of the samples in the box. Surprisingly, decontamination was more efficient when the samples was off-centered. This is completely surprising because samples positioned directly centered below the UV-LED ramp are expected to receive the maximum UV dose. Investigations revealed that an electrical transistor component had been welded in the wrong orientation, reducing the power of the UV-C LEDs used by a factor of 10, and triggering the random extinction of the central LED, explaining the results obtained. After correction, a new test campaign was then carried out on the boxes to validate the electrical modification. This time, complete disinfection up to the limit of detection was achieved, triggering a 4 Log10 (99,99%) reduction of infectious particle on both fabric and plastic, independently of the position of the sample in the device (Figure 2C). Altogether these results show that the use of ANCHOR viruses allow the rapid measurement of disinfection capacity of an UV irradiation device and can illustrate defects in disinfection efficiency. One can propose this kind of test to be used as quality control to ensure that the efficiency of commercial batch is in accordance with what was measured during laboratory testing or on a regular basis to verify disinfection efficiency over the life of the device. hAdv5 ANCHOR was used as a model to calibrate the device, and final validation was done on SARS-CoV-2 both using TCID50 and high content imaging. The device achieved 6log10 (99,9999%) reduction in SARS-Cov-2 infectious particles after 10 min (Figure 2E), triggering the complete disappearance of infected cells and viral RNA accumulation (Figure 2F). Disinfection using UV irradiation needs safety precautions as UV can trigger skin and eye damage. Also, the need for a power source can also limit its use. Other method such as passive decontamination can be useful to reduce virus burden in the environment.

**Figure 2.**
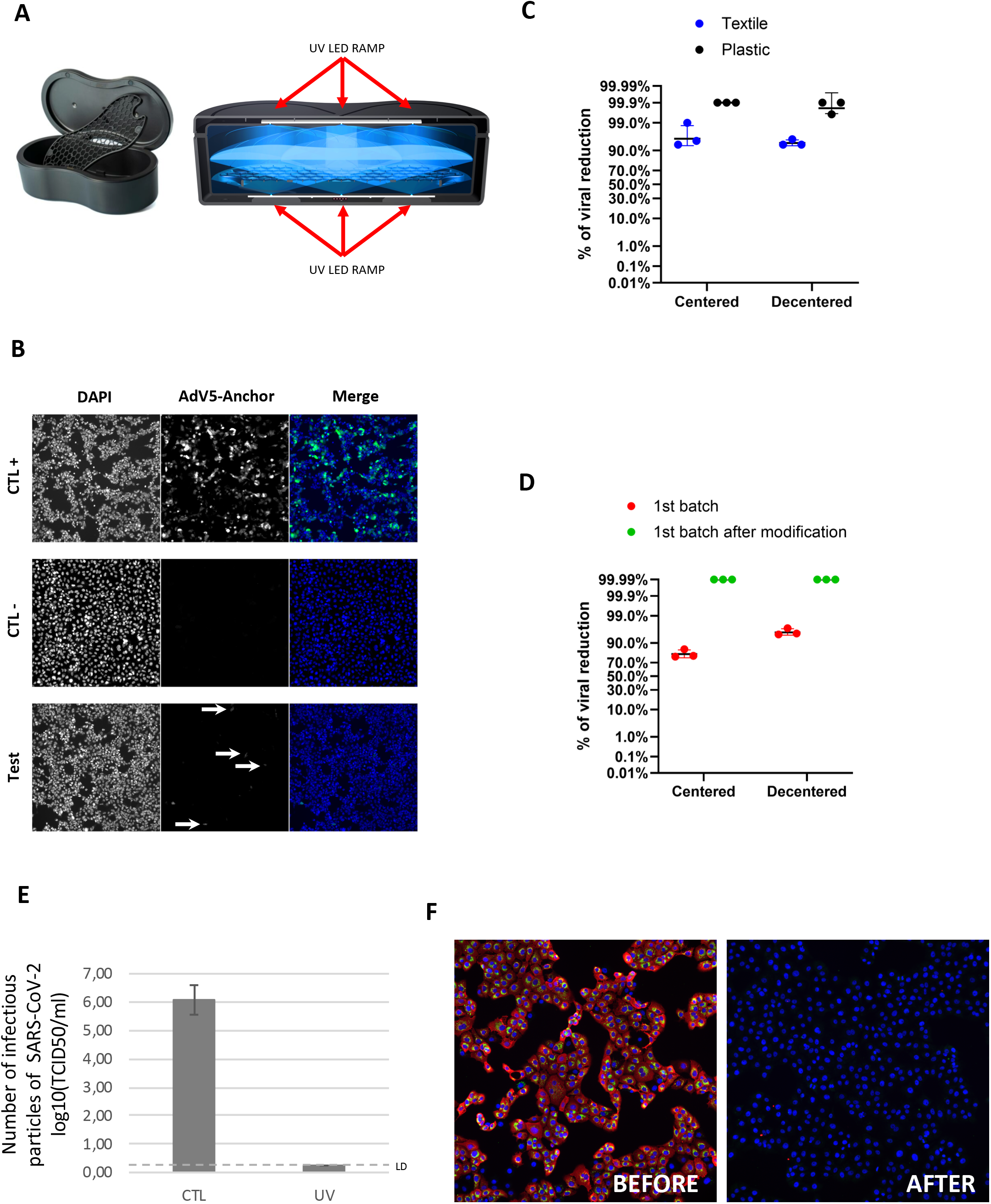
ANCHOR™ virus to develop and control UV disinfection solutions, from R&D to commercial product validation. **A:** Transversal section of the UVC disinfection box tested **B:** images illustrating the impact of UVC on the infectivity of the ANCHOR™ adenovirus. In green: cells infected by the adenovirus, in blue: cell nuclei. CTL+ is the viral load on the support but not irradiated /CTL-is the absence of virus in order to define the fluorescence threshold (background noise). **C:** quantification of the decrease in AdV5 ANCHOR™ infectious particles after irradiation, use of two different supports (fabric in blue and plastic in black). Results are expressed in per cent of infectious particles decrease compared to control unirradiated conditions. **D:** Monitoring production quality. Impact of a production defect on disinfection quality (in red), control after modification of the defective element (in green). Results are expressed in per cent of infectious particles decrease over unirradiated conditions. **E:** Efficacy testing on SARS-CoV-2. n=3, data expressed in log10 (TCID50) reduction **F:** Fluorescent labelling of cells infected with SARS-CoV-2 before and after UV irradiation in the box. Green: viral RNA, red: nucleocapsids, blue: cell nuclei. No detectable virus infection could be detected after irradiation.

### Use of ANCHOR viruses to investigate self-decontaminating surfaces activity

Passive decontamination can be achieved by using functional coatings such as pure or alloy of different metals as it is known for decade that metals ion such as copper can have anti-bacterial and antiviral activities. The problem underlying passive disinfection is the contact time required for achieving correct decrease in infectivity, which is usually several tens of minutes to hours. There is therefore a need to develop new alloys having fast and broad spectrum virus decay activities. We have tested new functionalized surfaces made by paint coating^18^. This coating can therefore be applied on a broad variety of surfaces such as metals, filters etc. Copper and therefore metal ions can act using multiple proposed mechanism of action, such as induction of oxygen radicals or alter lipid composition of the membrane, triggering desiccation and bacterial death. We therefore tested the activity of this new alloy, composed of different size copper and silver mixed particles, preventing copper oxidation and displaying prolonged activity. To verify if this new alloy was able to inactivate viruses, we first tested its activity on hAdv5 ANCHOR. Drops of viruses were laid under a microscopy slide to maximize contact with the functionalized surface for a 10min incubation. Compared to control conditions, functionalized surface shows 3 log10 (99,9%) decrease in hAdv particle infectivity, suggesting that the new coating is active on non-enveloped viruses (*data not show*). Testing the same alloy on VACV ANCHOR virus displayed strong reduction in VACV infectivity, triggering 2, 3 and >4log10 reduction in 5, 10 and 20min, respectively (Figure 3A). As shown in Figure 1, VACV is considered as a resistant virus and inactivation can be more rapid for other more sensitive enveloped viruses such as RNA viruses^15^. Indeed, testing the new alloy with extremely short contact time of 30 seconds and 1 min triggered the inactivation of above 90% of all enveloped RNA virus tested, namely influenza H1N1, highly pathogenous influenza H5N8 and SARS-CoV-2 (Figure 3B,C,D). Efficiency of the product was also verified on clinical Monkeypoxvirus strains circulating in Europe and triggered a 1.5 log10 decrease in 5 min and 4 log10 decrease after 10min on MPXV (data not shown). As more emergent viruses are expected to arise in the coming years, especially respiratory viruses and poxvirus such as MPXV, having a coating that can passively inactivate viruses within minutes will be of help to mitigate viral propagation. On another hand, this coating could be used in confinement laboratory, for example on benches or centrifuge, to reduce the environmental risk associated with pathogens manipulation especially in high confinement laboratories where leakage can lead to severe safety issues for the population. Also, with the increase number of BSL4 laboratories in the world, methods to control and ensure good disinfection are still needed. To our knowledge, few studies has shown BSL-4 suits disinfection efficiency, one using bacterial spore and the other one using an enveloped RNA virus^19,20^.

**Figure 3.**
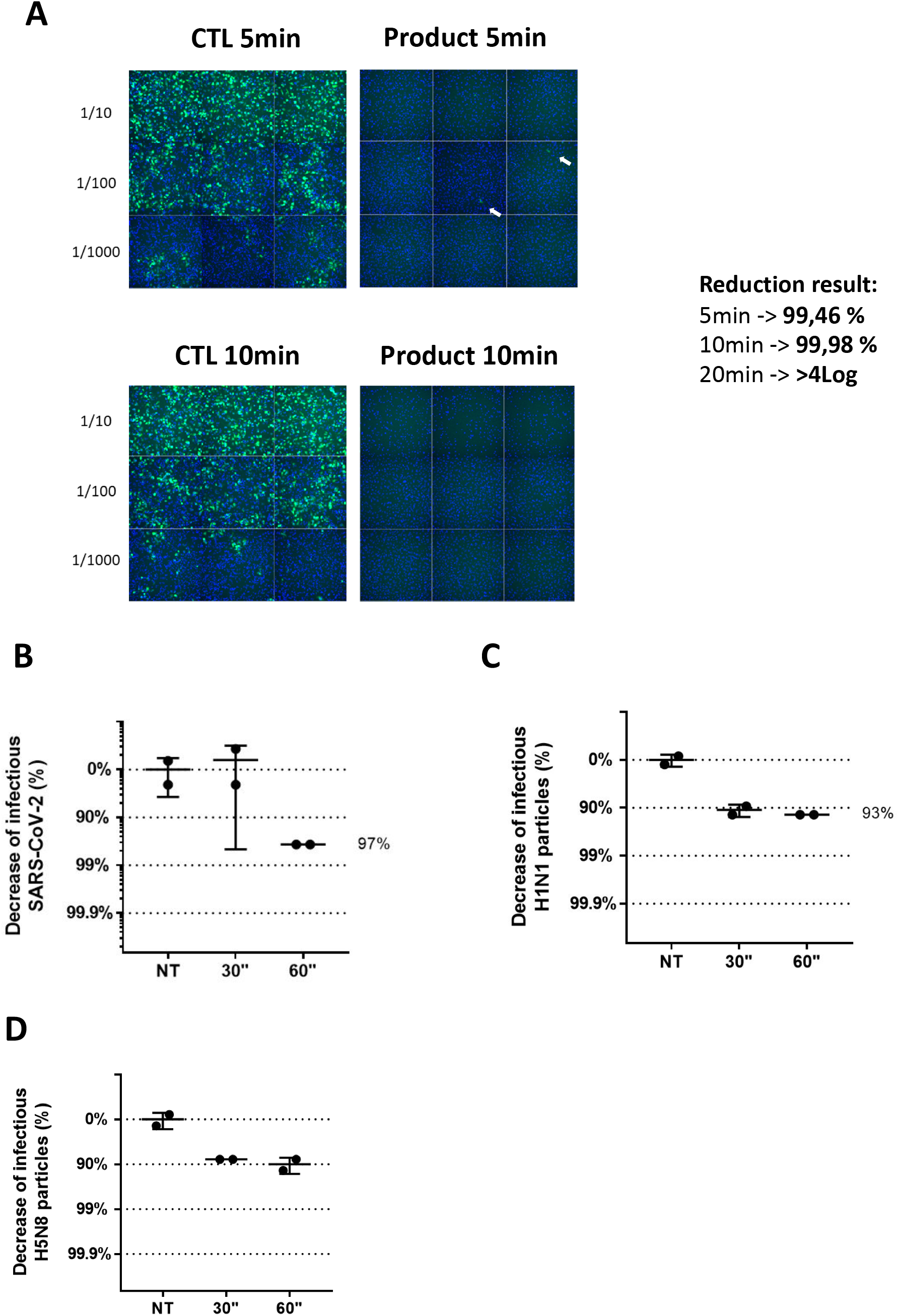
Analysis of the antiviral efficacy of self-decontaminating surfaces. ***A* :** Evaluation of the effectiveness of self-decontaminating surfaces on the VACV ANCHOR™ virus. Surfaces were contaminated with VACV ANCHOR™ (3,2.10^6^ ffu). Illustration of residual viral particles after contact with the surface. In green: VACV ANCHOR™, in blue: cell nuclei. **B:** Determination of the decontamination efficiency of surface on SARS-CoV-2. Surfaces were contaminated with a clinical isolate of SARS-CoV-2 (5.10^7^ TCID50; 25 µl of viral solution). Results are expressed in per cent of infectious SARS-CoV-2 decrease. **C**: Determination of the decontamination efficiency of surface on H1N1 influenza virus. Surfaces were contaminated with a H1N1 influenza virus strain (10^6^ TCID50; 25 µl of viral solution) Results are expressed in per cent of infectious H1N1 particles decrease. **D:** Determination of the decontamination efficiency of surface on H5N8 highly pathogenic influenza virus. Surfaces were contaminated with a H5N8 influenza virus strain (5.10^5^ TCID50; 25 µl of viral solution). Results are expressed in percentage of infectious H5N8 particles decrease.

### Use of ANCHOR viruses to evaluate disinfection efficiency on BSL4 suits and in operational BSL4 entry/exit conditions

Validating decontamination procedures in BSL4 laboratories is essential^20^. As our VACV ANCHOR virus is labeled “used by French Army forces”, we set up an evaluation of the BSL4 laboratory suit disinfection system to see if the processes used were efficient or could be optimized. We used both hAdv5 ANCHOR and VACV ANCHOR to artificially contaminate representative part of the pressurized suit (gloves, boots, full face shield) and the disinfectant evaluated under their current disinfection conditions (Bactipal 3% - 15min). As expected, results on AdV5 and VACV showed >5log10 viral load reduction (figure 4A-B), validating the disinfection conditions used by the laboratory on the specific BSL4 pressurized suits. It should be noted, however, that it was very difficult to recover the VACV virus from the boots.

**Figure 4.**
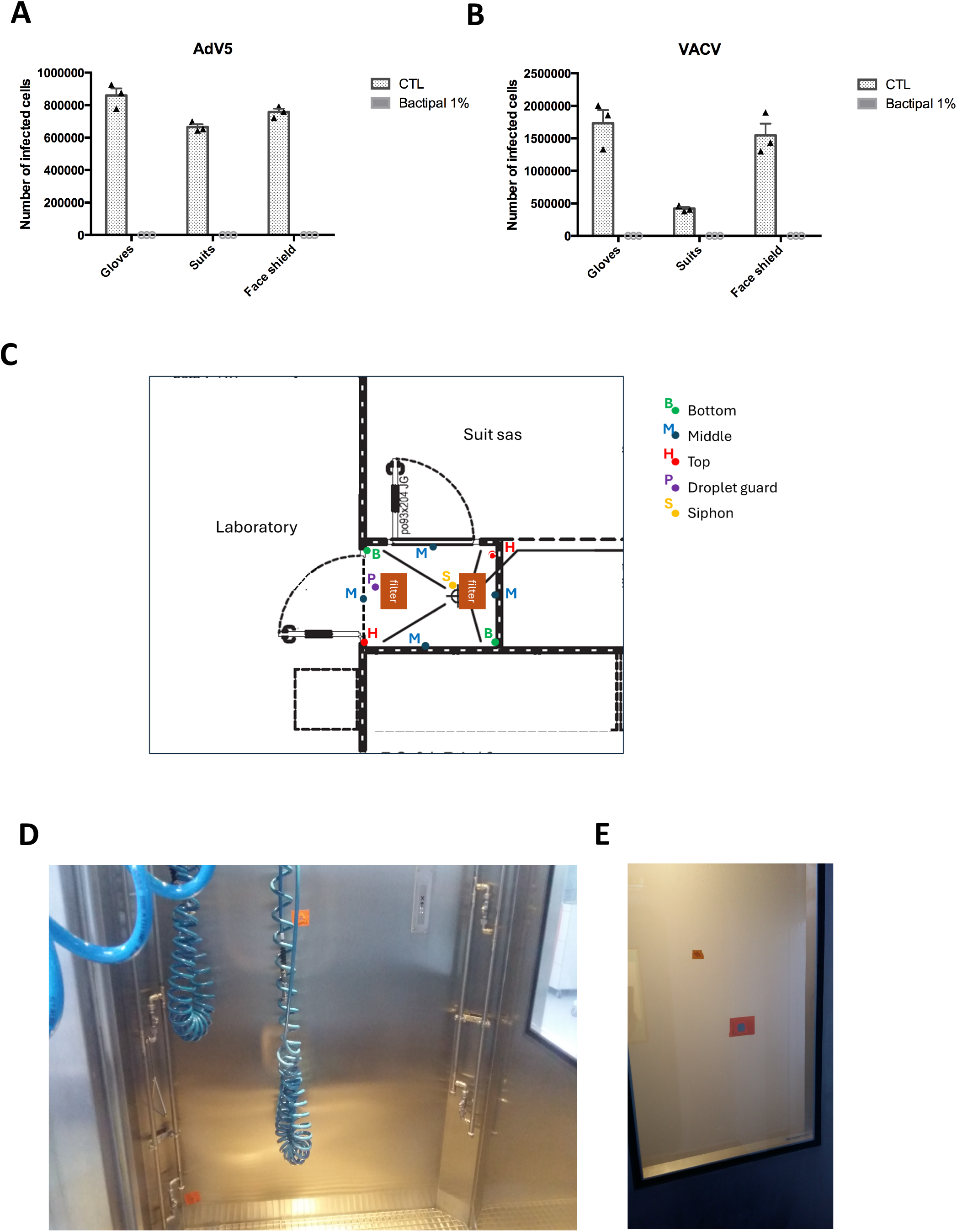
Application of ANCHOR technology to validate chemical disinfection in operational conditions. **A:** Efficiency test of the chemical decontamination protocol on BSL4 pressurized suits using AdV5 ANCHOR™ with or without chemical disinfection. **B:** Measurement of VACV ANCHOR™ particles with or without chemical disinfection. **C:** Layout plan of the biological VACV ANCHOR™ indicator carried out in the BSL4 showering room. Each dot indicates a zone of fixation of inox plates preloaded with VACV ANCHOR™. **D:** View of the inside of the cabin before spraying with decontaminant. **E:** View of the outside during fogging.

It seems that the more porous structure of the boots plastic “captures” the virus, which can cause problems during disinfection. Surprisingly, this result was specific to poxvirus, as no specific retention can be seen using hAdv5. These results could not have been obtained without testing the operational material with a live pathogen and highlighted the need to insist on boots disinfection. Therefore, contact time for the boots were extended to ensure that they were disinfected correctly, despite their porosity. Even if suits are correctly decontaminated, it may be different for the chemical shower room at the interface between the suits air lock and the interior of the BSL4 laboratory. To ensure that the showering room is evenly decontaminated, we use the VACV ANCHOR virus as a biological indicator for some of the qualification tests on the decontamination showers in the IRBA’s BSL-4 laboratory. Suspensions of the virus were deposited on stainless steel plates and allowed to dry before being installed in the shower cabins. (Figure 4 C-E). Then, showering was activated in nominal conditions. A reduction of 6log10 was demonstrated under the nominal operating conditions of the decontamination shower (Bactipal 1%, misting time of 4 min followed by 2 min rinsing with water). Therefore, VACV ANCHOR can be used as a biological indicator to validate disinfection efficiency in operational conditions and help mitigate risk for BSL-4 safety. Altogether, this project will lead to a better understanding and qualification of the efficiency of decontamination procedure and to identify and manage contamination risk. Use of mimetic autofluorescent virus allows rapid (overnight) evaluation of disinfection processes and could be used as a routine procedure to validate BSL-4 showering room decontamination and waste management for biohazard risk mitigation.

Altogether, we have shown that ANCHOR tagged viruses allow the rapid evaluation of radiative disinfection systems, physical decontamination systems and chemical disinfection. Finally, the imaging capabilities of our autofluorescent viruses are of significant asset for communication and marketing operations, allowing the direct visualization of the decontamination efficiency using an image based readout.

## Declaration of competing interest

FG is shareholder of NeoVirTech SAS. During this study, C.Q.F and EM were employees of NeoVirTech SAS. Other authors declare that they have no known competing financial interests.

### Author Contributions

Conceptualization: F.G. and E.M.; Experiments: F.G, E.M., V.P, C.Q.F, T.F, R.V, M.M, O.A, D.L, P.L, T.I, M.G. Supervision: F.G. and E.M.; Writing E.M– review & editing: F.G., C.Q.F and E.M. All authors have read and agreed to the published version of the manuscript.

## Materials and methods

### Cells

Human HEK293 (ATCC CRL-1573), Vero cells (ATCC CRL-1586) are grown in DMEM medium without phenol red (D1145; Sigma Aldrich) supplemented with 10% FBS (Eurobio-Scientific), 1mM sodium pyruvate (S8636; Sigma Aldrich), L-Glutamine (G7513; Sigma Aldrich) and Penicillin-Streptomycin solution (P0781; Sigma Aldrich).

### Viruses

VACV ANCHOR™: Vaccinia virus labelled with the ANCHOR system for real-time monitoring of viral infection (1.10^7^ ffu/mL). hAdv5 ANCHOR™: Human Adenovirus 5 labelled with the ANCHOR system (Viral titer 6.10^7^ffu/mL)^21^.hCMV ANCHOR™ Human cytomegalovirus labelled with ANCHOR system (1.10^8^ffu/mL). H1N1, H5N8 and SRAS-CoV-2 hCov19/France/OCC-IHAP-VIR12/2020 (see below). MPXV strain MPXV/France/IRBA2211/2022

### hAdv5 ANCHOR and VACV ANCHOR tests

HEK293 cells were seeded at 10^4^ cells/well in Corning Cellbind black glass-bottom 96-well plates in complete DMEM and infected 24 h post-seeding with ANCHOR engineered AdV5. For analysis, cells were directly stained with Hoechst 33342 (1 μg/ml) and imaged using a Thermo Scientific CellInsight CX7 microscope. Compartmental analysis was used to detect and quantify the number of ANCHOR-positive cells versus the total number of cells. For VACV, HeLa cells are plated at 10^4^ cells/well in Corning Cellbind 96 well plates in complete DMEM. 24 h post-seeding, surfaces or textiles were contaminated and incubated 15 min with viral solution. They were treated then at indicated time. Leftover contaminations were recovered by adding 500 µl of media. Cells received in triplicate 100µL. Cells are then incubated at 37 °C and 5% CO2 for 24h.Cells are fixed with 4% formalin (Sigma) for 10 min at RT, washed with PBS and incubated with PBS Hoechst 33342 (1mg/mL) and imaged using a Thermo Scientific CellInsight CX7 microscope. Compartmental analysis was used to detect and quantify the number of ANCHOR-positive cells versus the total number of cells.

### SARS-CoV-2 tests

In BioSafety Level 3 (BSL3) laboratory, surfaces or textiles were contaminated and incubated 15 min with a clinical isolate of SARS-CoV-2 (5.10^8^ TCID50; 25 µl of viral solution). They were treated then with UV-C at indicated time. Leftover contaminations were recovered by adding 500 µl of media. Four replicates were tested for each condition. Viral titers were determined by the TCID50 method on Vero-E6 cells and calculated by the Spearman & Karber algorithm. Surfaces were contaminated with a clinical isolate of SARS-CoV-2 (5.10^7^ TCID50; 25 µl of viral solution). Two replicates were tested for each condition: 30 and 60 sec of contact with the active surface and 60 sec of contact with the non-treated surface (NT). Leftover contaminations were recovered in 100µl of culture medium. For each sample, viral titers were determined by the TCID50 method on Vero-E6 cells and calculated by the Spearman & Kärber algorithm. Results are expressed in per cent of infectious SARS-CoV-2 decrease

### Determination of the decontamination efficiency of active surface on H1N1 and H5N8 influenza virus

Surfaces were contaminated with a H1N1 or H5N8 influenza virus strain (10^6^ TCID50 or 5.10^5^ TCID50 respectively; 25 µl of viral solution). Two replicates were tested for each condition: 30 and 60 sec of contact with the active surface and 60 sec of contact with the non-treated surface (NT). Leftover contaminations were recovered in 100µl of culture medium. For each sample, viral titers were determined by the TCID50 method on MDCK cells and calculated by the Spearman & Kärber algorithm. Results are expressed in per cent of infectious H1N1/H5N8 particules decrease.

### MPXV isolate

A primary MPXV isolate obtained from a patient during the 2022 French outbreak has been amplified on Vero-E6 cellls (ATCC) maintained in DMEM supplemented with 2.5% FCS. Briefly, viral stocks were prepared by infection of subconfluent Vero cells monolayers in 75 cm^2^ flasks with 5 ml of virion suspension adjusted at a M.O.I. of 0.01 in DMEM at 0.5% FCS.

After 90 min of incubation at 37°C under 5% CO2, the inoculum was removed and replaced by 20 ml of complete DMEM. Flasks were kept until cytopathic effect spread over at least 80% of the cell monolayer. Virions were extracted by three freeze-thaw cycles. Cell debris were removed by centrifugation 10 min at 3500 rpm. Viral suspension was aliquoted and kept frozen until use. Virus titer was determined on serial 10 fold dilutions inoculated on Vero cells monolayers seeded in 24 well plates using Karber method.

### Time dependant MPXV virus reduction on functionalized surfaces

Virus suspension was adjusted to 10^6^ PFU/100µl in DMEM without FCS. Spotted areas of 1.5 cm^2^ were marked. Drops of 100µl of virus suspension were loaded on functionalized surfaces and overlayed with a steril glass cover slide. At determined time points, virus was recovered by pipetting with an additionnal volume of 100µl to improve virus recovering. Remaining virus titers were determined by end point dilution.

### UV system

The UV system has been designed by Clinit technologie.

### Active surfaces

Patented by Lainisalo Capital OÜ

